# Nonlinear dynamics of chemotherapeutic resistance

**DOI:** 10.1101/300582

**Authors:** Y. Ma, P.K. Newton

## Abstract

We use a three-component replicator dynamical system with healthy cells, sensitive cells, and resistant cells, with a prisoner’s dilemma payoff matrix from evolutionary game theory to understand the phenomenon of competitive release, which is the main mechanism by which tumors develop chemotherapeutic resistance. By comparing the phase portraits of the system without therapy compared to continuous therapy above a certain threshold, we show that chemotherapeutic resistance develops if there are pre-exisiting resistance cells in the population. We examine the basin boundaries of attraction associated with the chemo-sensitive population and the chemo-resistant population for increasing values of chemo-concentrations and show their spiral intertwined structure. We also examine the fitness landscapes both with and without continuous therapy and show that with therapy, the average fitness as well as the fitness functions of each of the subpopulations initially increases, but eventually decreases monotonically as the resistant subpopulation saturates the tumor.

## I. INTRODUCTION

We describe an evolutionary game theory model of chemotherapeutic resistance [1] in a tumor with a pre-existing resistant cell. The model is based on a three-component replicator dynamical system with a frequency dependent fitness function based on a prisoner’s dilemma (PD) payoff matrix [2, 3]. Cell interactions occur between three cell types that form the tumor ecosystem: healthy cells (H); chemo-sensitive cells (S); and chemo-resistant cells (R). In a PD scenario, the healthy cells act as cooperators while the cancer cells act as defectors [4]. Unchecked, the defectors saturate the population as the average fitness decreases to a sub-optimal outcome. The goal of chemotherapeutics in this framework is to coax the defectors to cooperate, leading to a higher fitness Nash equilibrium [2, 3]. Growth rates are determined by cell fitness functionals which, in turn, depend on subpopulation sizes, i.e. they are frequency dependent.

The mechanism of resistance is based on the ecological notion of competitive release [5–7] of the resistant cell population when the sensitive cell population is reduced below a certain threshold. Above the threshold, the sensitive cells are able to outcompete (on average) the resistant cells due to the inherent *cost*-*of*-*resistance* which tilts the fitness landscape of the system in favor of the sensitive cell population allowing the tumor to grow. Under sufficient chemotherapeutic pressure, the sensitive cell population is reduced enough to allow the resistant population to begin to flourish and eventually regrow the tumor in a form that is much harder to treat. A quantitative understanding of this phenomena is necessary in order to develop chemotherapeutic strategies (i.e. adaptive therapies) to combat it, a point of view adopted and developed in [8–13].

## II. THREE-COMPONENT REPLICATOR SYSTEM

The model we develop is based on a three-component replicator nonlinear dynamical system governing three competing subpopulations of cells: 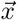 = (*x*_1_,*x*_2_,*x*_3_)^*T*^ = (*x_H_*, *x_S_*, *x_R_*)^*T*^, where *x*_1_ represents the proportion of healthy cells (H), *x*_2_ represents the proportion of sensitive cells (S), and *x*_3_ represents the proportion of resistant cells (R), with *x*_1_ + *x*_2_ + *x*_3_ = 1. The equations which describe their interaction are:

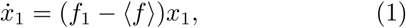

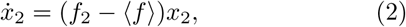

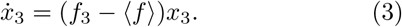

with *f_i_* representing the fitness of each of the subpopulations (*i* = 1, 2, 3) as their relative populations change, and 〈*f*〉 representing the average fitness of the entire population. The exponential growth-decay rates of each of the subpopulations are then determined by (*f_i_* − 〈*f*〉) which dictates whether the subpopulation fitness is above or below the average population fitness, hence whether the subpopulation decays or grows.

The fitnesses of the three subpopulations are defined by the linear functionals:

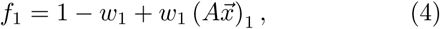

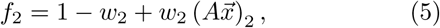

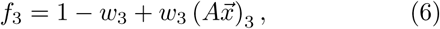

where 0 ≤ *w_i_* ≤ 1, (*i* = 1, 2, 3) are selection pressure parameters we use to shape the fitness landscapes. The average population fitness is defined by the quadratic functional:

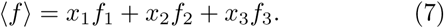

It is straightforward to see that the fixed points of the system (1)-(3) are of three basic types: (i) There are the three fixed points at each of the corners of the triangular phase space diagram shown in Figure 1, when two of the subpopulation values are zero, and the third is saturated: (*x*_1_, *x*_2_, *x*_3_) = (*x_H_*, *x_S_*, *x_R_*) = (1, 0, 0); (0, 1, 0); (0, 0, 1); (ii) There are three possible fixed points on the triangle sides, which correspond to one of the subpopulation values equaling zero, with the other two having fitness values equal to the population average: *x*_1_ = 0, *f*_2_ = 〈*f*〉, *f*_3_ = 〈*f*〉; *x*_2_ = 0, *f*_1_ = 〈*f*〉, *f*_3_ = 〈*f*〉; *x*_3_ = 0, *f*_1_ = 〈*f*〉, *f*_2_ = 〈*f*〉; (iii) There is the *balanced fitness state*, when none of the subpopulation values is zero, but each of the subpopulation fitness values equals the population average: *f*_1_ = 〈*f*〉; *f*_2_ = 〈*f*〉; *f*_3_ = 〈*f*〉. Which of these fixed points lies on or inside the triangular phase space, and their stability properties, depend in detail on the parametric values, which we describe in §III.

**FIG. 1.**
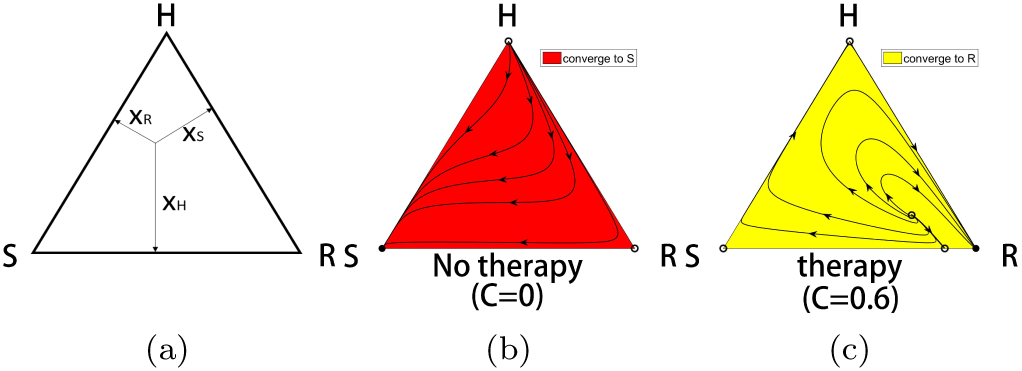
The competitive release mechanism. (a) The three-component phase space associated with competing populations of (*H*, *S*, *R*) cells. (b) With no therapy, the *S* corner of the triangle is a globally attracting fixed point, while the *H* and *R* corners are unstable. All initial conditions lead to a saturated tumor. Filled circles are stable, unfilled circles are unstable. (c) For continuous chemotherapy above a threshold level, the *R* corner of the triangle is a globally attracting fixed point, while the *H* and *S* corners are unstable. All initial conditions (except those lying on the separatrix connecting the interior balanced fitness state to the *S* – *R* side) lead to a resistant tumor.

### A. The prisoner’s dilemma as a cancer model

The standard version of the PD payoff matrix [2] in a 2 × 2 setting in which healthy cells compete with cancer cells is:

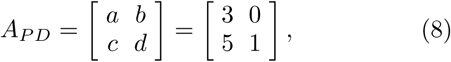

where the first row and column correspond to the payoffs associated with the *cooperator* (C) in the PD evolutionary game, and the second row and column correspond to the payoffs associated with the *defector* (D). In the simplest tumor growth paradigm in which a population of healthy cells competes with a population of cancer cells, the healthy cells are the cooperators, while the cancer cells are the defectors. In any mixed population 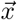 = (*x_C_*,*x_D_*)^*T*^, 0 ≤ *x_C_* ≤ 1; 0 ≤ *x_D_* ≤ 1; *x_C_* + *x_D_* = 1, the fitness functions, 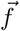 = (*f_C_*, *f_D_*)^*T*^, associated with the two subpopulations are:

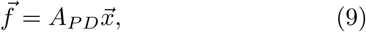

which in component form yields:

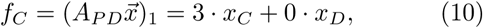

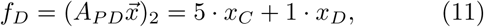

while the average fitness of the total population is given by the quadratic form:

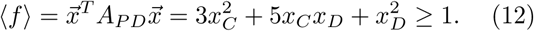

Note that the average fitness of the healthy state (*x_C_*,*x_D_*) = (1,0) is given by 〈*f*〉_*C*_ = 3, while that of the cancerous state (*x_C_*,*x_D_*) = (0,1) is given by 〈*f*〉_*D*_ = 1, which minimizes the average fitness. Tumor growth is then modeled as a 2 × 2 evolutionary game governed by the replicator dynamical system:

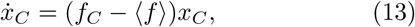

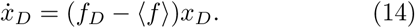

It is straightforward to show:

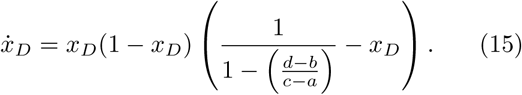

From this, we can conclude that for any initial condition containing at least one cancer cell: 0 < *x_D_* (0) ≤ 1, we have:

i. *x_D_* → 1, *x_C_* → 0 as *t* → ∞
ii. 〈*f*〉 → 1 as *t* ↑ ∞.

Condition (i) guarantees that the cancer cell population will saturate, while condition (ii) guarantees that the saturated state is sub-optimal, since 〈*f*〉_*D*_ < 〈*f*〉_*C*_. For these two reasons, the prisoner’s dilemma evolutionary game serves as a simple model for tumor growth both in finite population models, as well as replicator system (infinite population) models [14–17].

Generalizing to a three-component system, the fitness functions (4)-(6) are defined via a payoff matrix *A* which is a 3 × 3 matrix defining the evolutionary game played by each of the cells on pairwise interactions. For this, we take every 2 × 2 sub-matrix to be a PD game, i.e. we take A to be the 3 × 3 PD matrix:

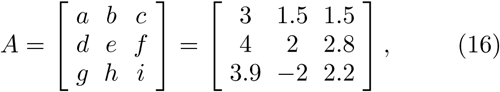

with the PD inequalities [2]:

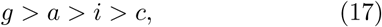

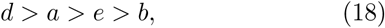

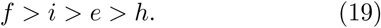

The numerical values in (16) are chosen for convenience and satisfy the constraints (17)-(19). In each cell-cell interaction, the healthy cells *x*_1_ (healthy H) are cooperators, and the two-species of cancer cells, *x*_2_ (sensitive S), and *x*_3_ (resistant R), are the defectors. In any interaction between a chemo-sensitive cell (S) and a chemo-resistant cell (R), the sensitive cell is the defector, while the resistant cell is the cooperator. The payoff matrix (16) guarantees that for any interaction between two cells, the system retains features (i),(ii) detailed previously (i.e. tumor growth leading to sub-optimal fitness). In addition, the payoff matrix also imposes a *cost to resistance* if we add the extra constraint *d* > *g* which guarantees:

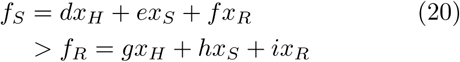

It is worth pointing out that shaping the fitness landscape by adjusting the selection parameters (*w*_1_,*w*_2_,*w*_3_) is equivalent to choosing an adjusted payoff matrix *A* in the fitness equations (4)-(6) so that:

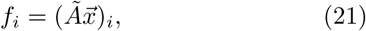

where:

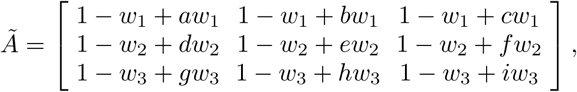

To see this, use the fact that (1 − *w_i_*) ≡ (1 − *w_i_*)(*x*_1_ + *x*_2_ + *x*_3_) in the fitness equations (4)-(6).

### B. Chemotherapy model

To model chemotherapeutic response, we exploit the fact that chemotherapy alters the fitness landscape of the cell populations, decreasing the fitness of the sensitive population to a level below that of the resistant population [18]. We introduce a chemo-concentration parameter *C* (dose concentration per unit time) which acts on the sensitive cell population (linearly):

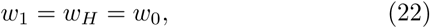

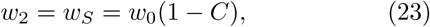

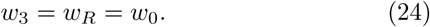

The total dose administered is *D* = *C* · *t*, where *t* is the time administered. Here *w*_0_ sets the timescale in the replicator system (which we normalize to *w*_0_ = 1) to obtain:

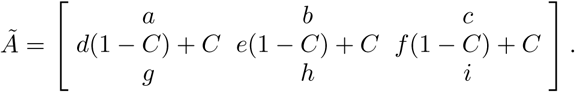

The concentration parameter *C* adjusts the selection pressure on the sensitive cell population by adjusting the entries of the payoff matrix (linearly) thereby shifting the relative fitnesses of the subpopulations even to the point that *Ã* is no longer a PD matrix as it was for *C* = 0. We examine properties of the phase space of the dynamical system for constant values of *C* (continuous therapy) for values 0 ≤ *C* ≤ 1.

## III. RESULTS

Figure 1 shows the mechanism of chemotherapeutic resistance via competitive release determined by the system (1)-(3), as depicted in the triangular phase space diagram shown in Fig 1(a). With no therapy (Fig.1(b), *C* = 0), the sensitive corner *S* is globally attracting, while the *H* and *R* corners are unstable. When the continuous therapy parameter *C* = 0.6 (Figure 1(c)), the resistant corner is globally attracting, while the *H* and *S* corners are unstable. Filled corners are stable, unfilled corners are unstable. This is the basic mechanism of competitive release induced by sufficiently strong chemotherapeutic dose [19].

Intermediate values of *C* reveal a much more complex picture. We show in Fig 2 the location of the fixed points as a function of *C*. For values 79/228 = 0.34649… < *C* < 0.7, the balanced fixed point state is an interior fixed point, which forms the central spiral associated with the basin boundaries between the asymptotically stable *S* corner and *R* corner, shown in the panel in Figure 3. Also shown in the figure are the nullclines defined by the curves *dx_H_/dt* = 0, *dx_S_/dt* = 0, *dx_R_/dt* = 0. Opposite sides of each of the nullclines mark a change in whether the particular subpopulation decays or grows. Stable fixed points are filled circles, unstable are unfilled circles. The mixed population state exists between values 1/3 < *C* < 1/2 where the basin of attraction sizes (areas), shown in Figure 4, change rapidly (as *C* varies), one at the expense of the other, in an intertwined spiral structure centered at the balanced fixed point state. The rapid transition between the two states occuring for small changes in the chemo-concentration parameter highlights the sensitivity of the system to chemotherapeutic dosing levels. The fitness landscapes are shown in Figure 5 both for *C* = 0 and *C* = 0.6. With no therapy, the fitness curves are monotonically decreasing functions as the sensitive population saturates the tumor in a sigmoidal shaped [14] growth curve. With continuous therapy above threshold, the fitness curves initially increase (i.e. tumor regression), but eventually decrease monotonically as the tumor relapses. The healthy cell population initially increases, but eventually the resistant population saturates the tumor.

**FIG. 2.**
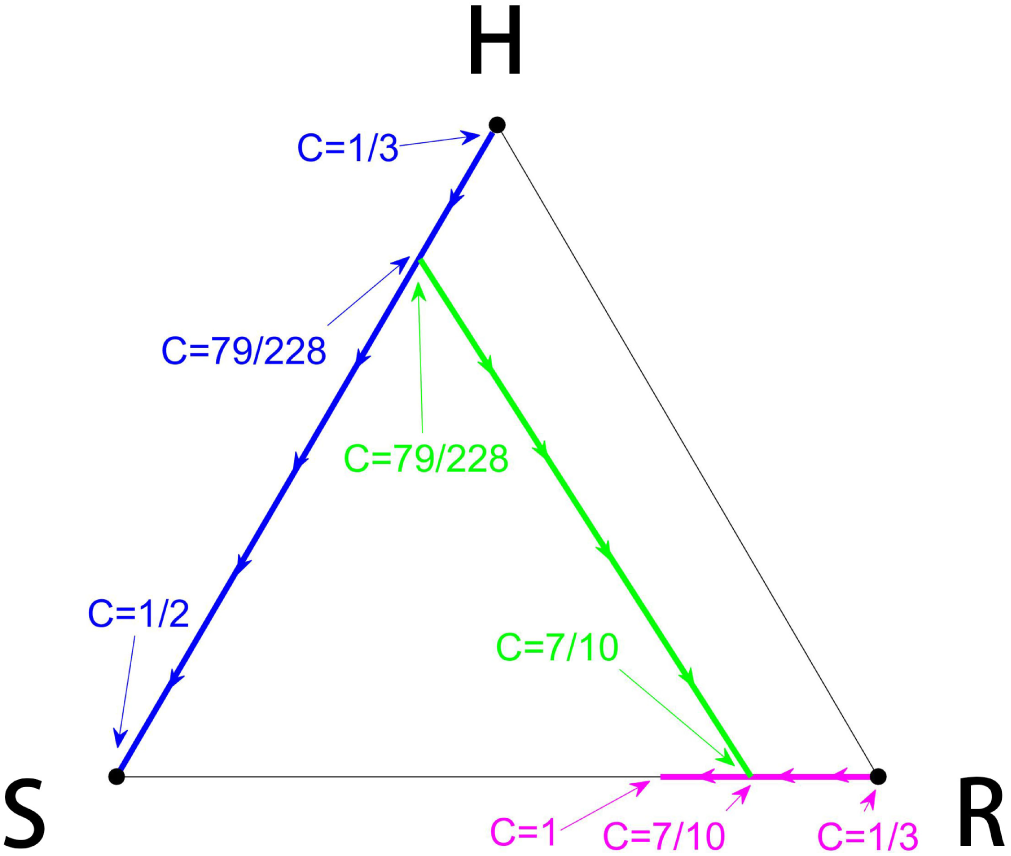
Fixed point locations. Tracking the location of the fixed points as a function of chemo-concentration parameter *C*. The fractional values indicate that the values are analytically obtained.

**FIG. 3.**
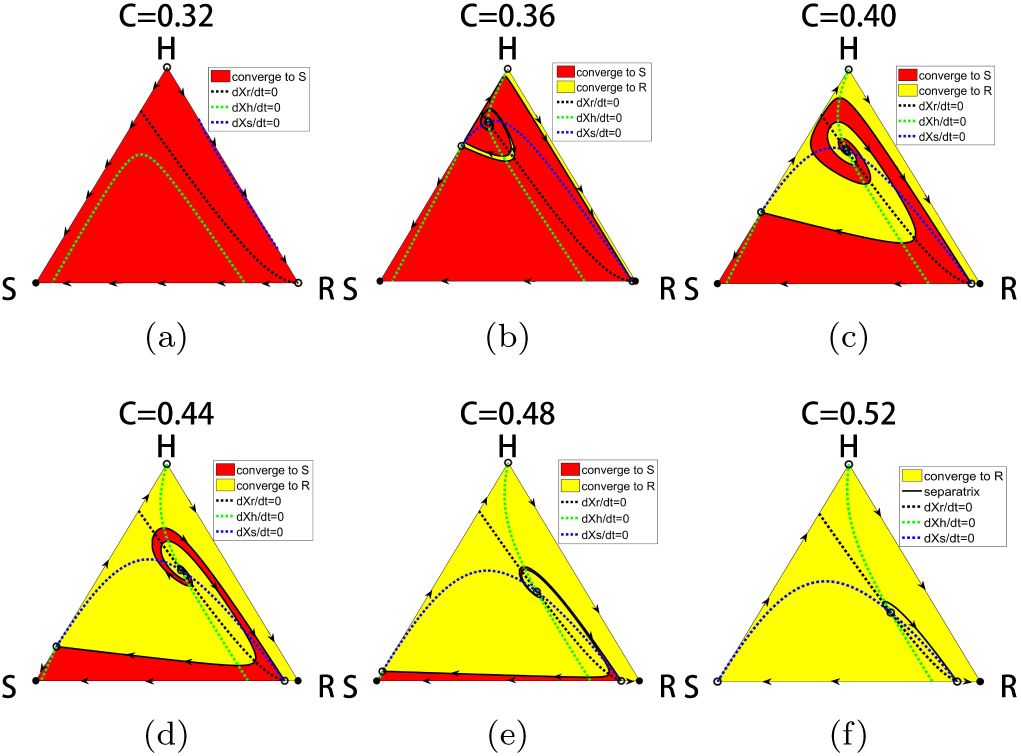
Basins of attraction of *S* and *R* states. Panel showing the separatrices through the balanced fitness interior fixed point that determines the basin boundaries of attraction of the *S* state and the *R* states. The interior fixed point is the one in which each of the sub-population fitness levels exactly matches the average fitness of the entire population. Filled circles are stable, unfilled circles are unstable.

**FIG. 4.**
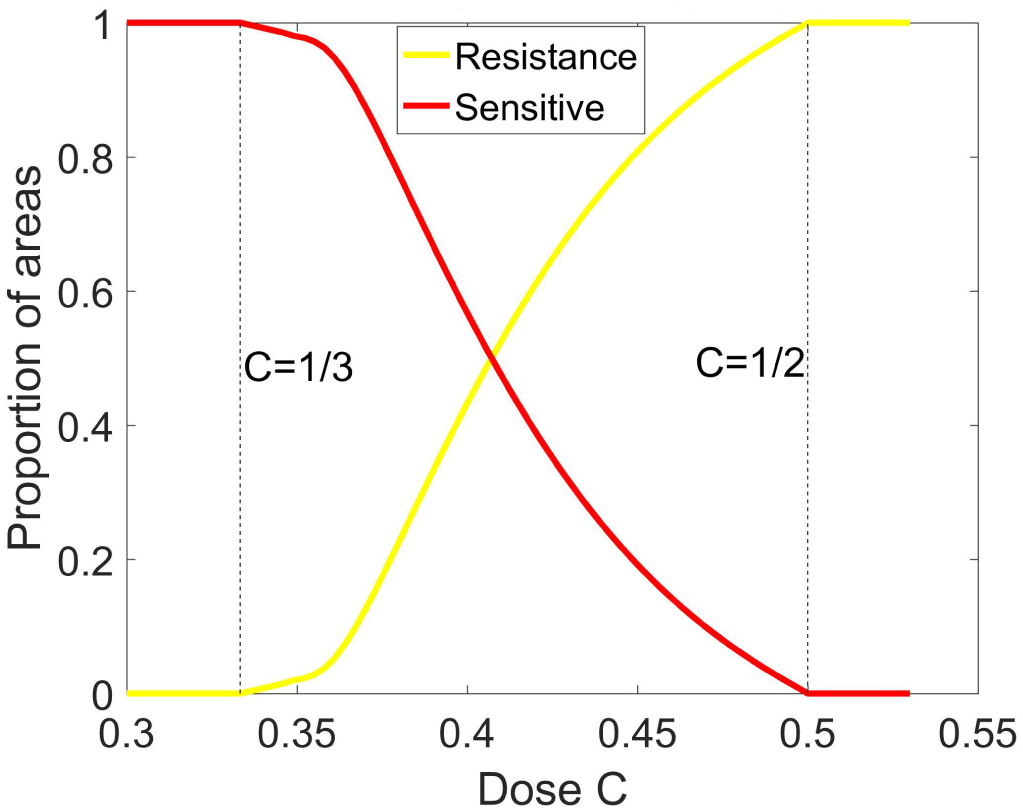
Basin of attraction areas. Areas of basin of attraction of *S* fixed point and *R* fixed point as a function of chemo-concentration *C*.

**FIG. 5.**
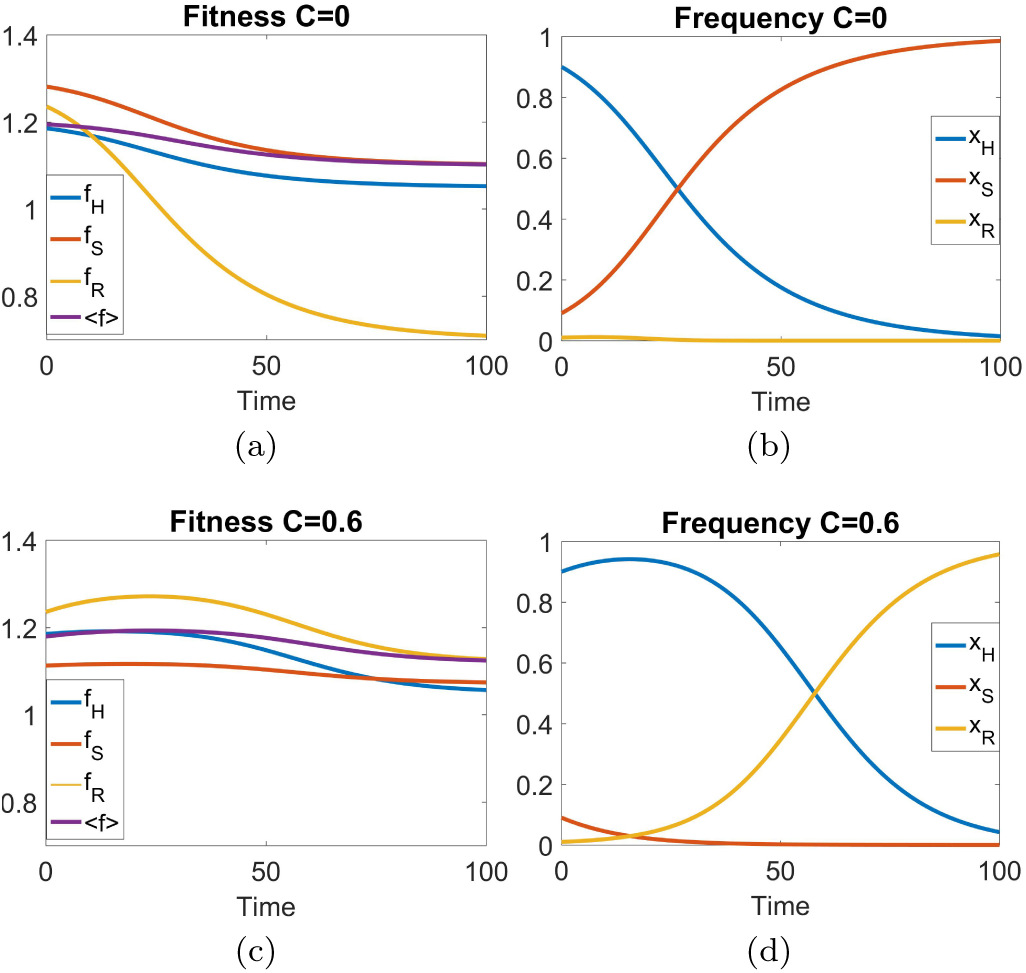
Fitness landscapes as a function of time. With no therapy (*C* = 0), the fitness curves are contunuously decreasing functions as the tumor saturates with the sensitive cell population in a sigmoidal shaped growth curve. With continuous therapy (*C* = 0.6), the fitnesses initially increase indicating tumor regression, but eventually decrease. The healthy subpopulation initially increases before the resistant population eventually saturates the tumor. Initial condition (0.9,0.09,0.01) with *w*_0_ = 0.1

## IV. DISCUSSION

The model shows that if the chemo-dose exceeds a threshold value of *C* > 0.5, the tumor may regress for a period of time, but eventually regrows to form a resistant tumor (Figure 1(c)) as long as there is at least one resistant cell in the population. This process of competitive release is very robust and occurs for all initial distributions of the three subpopulations. For chemo-doses that are not as large, the results are much more sensitive to small changes in *C* since the phase space diagram generally has two basins of attraction, one associated with a saturated sensitive state, the other with a saturated resistant state, and these basins of attraction are spirally intertwined. Small differences in the balances among the three types of cells comprising our model can determine whether the long-time dynamics converges to the *S* state or the *R* state. The relative areas of the basins of attraction also sharply change with small changes in chemo-dose values between 1/3 < *C* < 1/2 reflecting the sensitivity in choosing the optimal chemo-dose levels. There is no value of *C* for which the system converges to the *H* state, as this state is always unstable. However, because of the detailed and intertwined properties of the phase space basins of attraction for different values of *C*, one could imagine exploiting this phase space structure using time-dependent chemo-schedules (i.e. *C*(*t*)) to maintain sub-population balances *near* the H corner for long enough periods of time to effectively *manage and control* the growth of the tumor without necessarily eliminating it [8, 9, 20]. In fact, adaptive therapies where chemo-schedules are altered during the course of treatment based on tumor response, are currently being clinically tested [21] and show promise. Using C(t) as a control accuator for the replicator system (1)-(3) in a closed-loop adaptive control scheme will be described in a separate publication.

## Acknowledgments

We gratefully acknowledge partial support from the Breast Cancer Research Foundation (BCRF) and the Jayne Koskinas & Ted Giovanis Foundation (JKTG) for Health and Policy. We thank S. Anderson, D. Basanta, J. Brown, H. Enderling, R. Gatenby, and J. West at the Moffitt Cancer Research Center for helpful conversations regarding this and related work.

